# A targetable opioid/cancer associated fibroblast axis drives extracellular matrix remodeling and tumor aggressiveness in pancreatic cancer

**DOI:** 10.64898/2026.05.20.726559

**Authors:** Kathryn E. Maraszek, Arwen A. Tisdale, Hunter D. Reavis, Xiaozhuo Liu, Eduardo Cortes Gomez, Aleksandr Dolskii, Caneta Brown, Daksh Thakkar, Maura Dungan, Anusha Adhikari, Emily Mackey, Janusz Franco-Barraza, Brooke A. Pereira, Paul Timpson, Dean G. Tang, Nina Steele, Edna Cukierman, Michael E. Feigin

## Abstract

Pancreatic ductal adenocarcinoma (PDAC) is an intractable disease with few effective treatment options. PDAC is characterized by a dense, fibro-inflammatory tumor microenvironment (TME) consisting mainly of cancer-associated fibroblasts (CAFs) and a CAF-generated collagen-rich extracellular matrix (ECM). As the ECM has profound impacts on tumor progression and therapy response, it is critical that we understand the mechanisms underlying ECM deposition and remodeling. In addition to a highly fibrotic and reactive TME, a hallmark of PDAC is pain. 93% of PDAC patients experience pain, and ∼70% are prescribed opioids for pain management during the course of their cancer treatment. Despite epidemiological evidence linking opioid use with diminished patient survival, how opioids impact tumor biology remains largely unknown. We now provide evidence that both endogenous and exogenous opioids drive ECM remodeling in the PDAC TME. We find that the commonly prescribed opioid morphine promotes the development of poorly differentiated tumors and increases collagen bundling and maturation in a mouse model of PDAC. Accordingly, RNA sequencing reveals that morphine induces significant upregulation of ECM genes and collagen modifying enzymes. We developed a morphine-induced gene signature which correlates significantly with the basal/mesenchymal subtypes of human PDAC and predicts worse overall survival in PDAC and other tumor types. Mechanistically, pharmacological inhibition and genetic knockdown of the mu opioid receptor (OPRM1) in CAFs attenuates expression of type 1a and type 3a collagens, and the myofibroblastic CAF marker alpha-SMA, demonstrating that opioid signaling is a direct regulator of CAF biology. Additionally, we provide the first evidence that CAFs produce endogenous opioids capable of activating OPRM1 and driving collagen expression. Finally, treatment with the FDA-approved peripherally restricted OPRM1 antagonist methylnaltrexone (MNTX) reduces desmoplasia, tumor weight, and ascites burden in a mouse model of PDAC. Therefore, we have identified a novel opioid-mediated signaling axis driving PDAC desmoplasia and reveal MNTX as a potential therapeutic to inhibit both exogenous and endogenous opioid-induced ECM remodeling and tumor aggressiveness.

## Introduction

Pancreatic ductal adenocarcinoma (PDAC) is a highly lethal malignancy with a dismal five-year survival rate of 13.3% [1, 2]. Nearly 50% of patients are diagnosed at an advanced, metastatic disease stage, at which the five-year survival rate drops to 3.2% [1, 3]. Although PDAC only accounts for 3% of newly diagnosed cancer cases in the US, it is currently the third-leading cause of cancer-related mortality and is expected to be the second-leading cause by 2030 [4, 5]. Surgical resection continues to be the only potentially curative treatment option; however, only 10-15% of patients are eligible for surgery at the time of diagnosis, with the vast majority presenting with either locally advanced or metastatic disease [3].

PDAC is characterized by a dense, desmoplastic stroma, which can comprise the majority of the tumor volume. This unique stroma is made up predominantly of functional interstitial units made up of cancer-associated fibroblasts (CAFs) and the extracellular matrix (ECM) proteins that they produce, such as collagens [6]. The production and remodeling of the PDAC ECM by CAFs regulates disease progression and therapeutic efficacy [7]. Excessive deposition of ECM proteins impedes vascular function and increases interstitial fluid pressure within the tumor microenvironment (TME), hindering drug delivery and promoting chemoresistance [8]. Additionally, CAF/ECM units and the ECM can drive tumor progression through providing nutrients that foster tumor cell proliferation and survival [8–11]. The structure of the ECM is also of importance, as increased stiffness due to collagen crosslinking and bundling can promote tumor cell proliferation, epithelial-to-mesenchymal transition, and a reduction in tissue polarity [12–15]. Similarly, increased collagen fiber thickness has been associated with worsened PDAC survival [16]. Areas of increased thickness and alignment of collagen fibers correspond to sites of tumor invasion, and correlate with worse survival in breast and other cancers [17]. Thus, understanding factors that contribute to the deposition and remodeling of the ECM is crucial in the effort to improve therapeutic efficacy and treatment outcomes for patients with PDAC.

A hallmark of PDAC is the patient experience of pain. 93% of patients report that they experience pain, with 83% ranking their pain as moderate-to-severe [18]. As a result, approximately 75% of PDAC patients receive prescription opioids across all stages of disease [19]. Importantly, both opioid usage and expression of the mu opioid receptor (OPRM1), the predominant receptor through which opioids signal, correlate with worse survival in various cancer types, including PDAC [20, 21]. While previous studies have addressed the direct effects of opioid signaling on tumor cells and the immune component of the TME, there are currently no published studies on the effects of opioids on CAFs or the ECM [22–24].

In this study, we report a novel, targetable mechanism for exogenous and endogenous opioids in driving pancreatic tumor aggressiveness and regulation of CAF-mediated ECM remodeling. Using both *in vivo* and *in vitro* models, we find that exogenous opioids promote collagen expression and pro-tumorigenic ECM remodeling. We develop a morphine/ECM gene signature that correlates with poor outcome in PDAC and other tumor types, highlighting the potential for this axis to be conserved across the cancer landscape. We also report the discovery that CAFs produce endogenous opioids that drive collagen production, even in the absence of exogenous prescription opioids. Critically, we find that inhibition of peripheral opioid signaling using the peripherally restricted OPRM1 antagonist methylnaltrexone (MNTX) reduces tumor aggressiveness *in vivo*. To our knowledge, this is the first study to demonstrate that endogenous and exogenous opioids drive ECM remodeling in the TME, providing a novel mechanism of ECM regulation. Furthermore, we provide evidence that MNTX, an FDA-approved drug already in use in cancer patients, may represent a novel therapeutic to inhibit opioid-induced aggressiveness in PDAC.

## Results

### Morphine promotes poor differentiation and alters collagen structure in the pancreatic tumor microenvironment

To determine if opioids impact murine PDAC biology, we orthotopically implanted LSL-KrasG12D/+; LSL-Trp53R172H/+; Pdx-1-Cre (KPC) tumor pieces into the pancreata of wild-type C57BL/6 mice, allowed the tumors to establish for three days, then treated the mice with morphine or vehicle control for two weeks **(Fig. 1A)**. Morphine was chosen for this study as it is among the most commonly prescribed opioids to PDAC patients and the most well-characterized experimentally in mouse models. Morphine dose and administration were chosen based on published reports demonstrating therapeutic effects on pain mitigation in mice [25]. Although tumor size did not differ significantly upon morphine treatment **(Supp Fig. 1A)**, histopathology revealed pronounced differences in both tumor epithelium and stroma. Notably, tumors from vehicle-treated mice were predominantly well-to-moderately differentiated, while tumors from morphine-treated mice were nearly all (87%) poorly differentiated **(Fig. 1B, C)**. Tumors from morphine-treated mice displayed increased positivity for Ki67, indicating increased proliferative capabilities **(Fig. 1D)**. We also noted an increase in the myofibroblastic CAF marker alpha smooth muscle actin (αSMA) in tumors from morphine-treated mice **(Fig. 1E)**, suggesting alterations to the tumor stroma. As CAFs are the primary producers of collagen in the PDAC ECM, we assessed collagen deposition by Masson’s trichrome and picrosirius red staining. Tumors from morphine-treated mice exhibited notable increased staining for collagen through both methods **(Fig. 1F)**. To assess the polymerization status of the increased collagen within the stroma, we used multiphoton second harmonic generation (SHG) microscopy, by which polymerized collagen fibers and their architectural features can be evaluated. There were no statistically significant changes in collagen fiber alignment or length **(Supp Fig. 1B, C)**. However, this approach revealed a statistically significant opioid-induced increase in collagen fiber size and density **(Fig. 1G, H)**, indicating increased collagen bundling and maturation, features associated with PDAC progression and poor patient outcomes. Overall, these results demonstrate that morphine alters the entire PDAC tumor mass, including both epithelial and stromal compartments.

**Figure 1:**
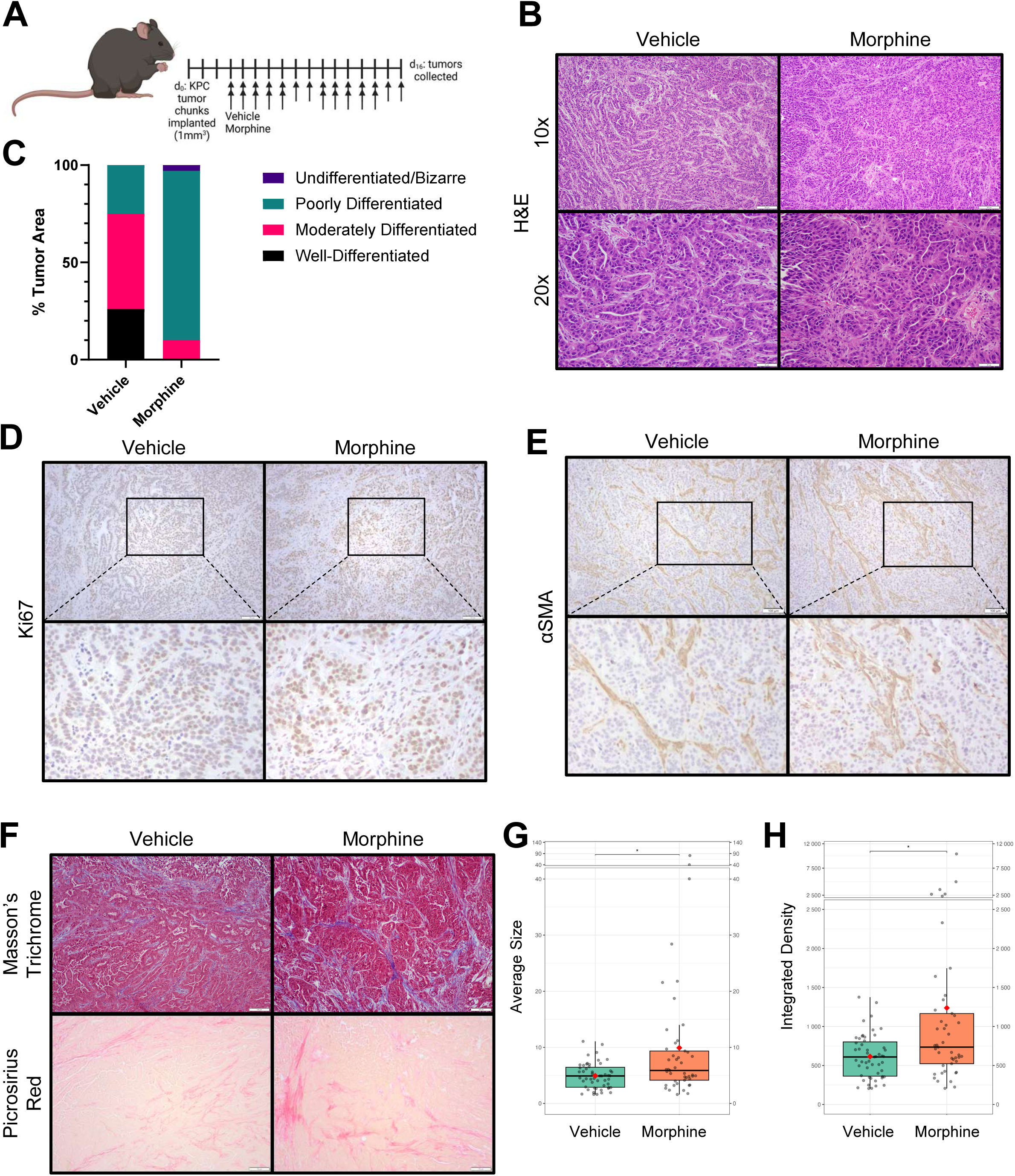
Morphine alters tumor phenotype and collagen structure in the pancreatic tumor microenvironment. **A,** Experimental schematic of orthotopic transplantation of LSL-KrasG12D/+; LSL-Trp53R172H/+; Pdx-1-Cre (KPC) tumors into C57BL/6 mice and treatment with subcutaneous vehicle/morphine at 10mg/kg (*n =* 10/arm) for 14 days. Arrows indicate doses on each individual day. **B,** Representative 10x (top) and 20x (bottom) H&E images of tumors from vehicle (left) and morphine-treated (right) mice. Scale bars represent 100µm (top) and 50µm (bottom). **C,** Quantification of total tumor area (%) by differentiation status between vehicle and morphine-treated mice. **D,** Representative 10x (top) images of Ki67 IHC in tumors from vehicle (left) and morphine-treated (right) mice with zoomed in insets (bottom). Scale bars represent 100µm. **E,** Representative 10x (top) images of αSMA IHC in tumors from vehicle (left) and morphine-treated (right) mice with zoomed in insets (bottom). Scale bars represent 100µm. **F,** Representative 10x images of Masson’s Trichrome (top) and Picrosirius Red (bottom) staining of tumors from vehicle (left) and morphine-treated (right) mice. Scale bars represent 100µm. **G,** Quantification of collagen fiber size by SHG microscopy in tumors from vehicle and morphine-treated mice. **H,** Quantification of collagen fiber density by SHG microscopy in tumors from vehicle and morphine-treated mice. Statistical comparisons were made using the Wilcoxon test. *, *p* < 0.05; **, *p* < 0.01; ***, *p* < 0.001; **** *p* < 0.0001.

### Morphine induces expression of an extracellular matrix signature associated with poor survival

To investigate changes in gene expression associated with morphine treatment, RNA sequencing was performed on tumors harvested from vehicle and morphine-treated mice **(Fig. 2A)**. RNA sequencing of tumors from 5 vehicle-treated and 5 morphine-treated mice revealed 362 and 264 gene transcripts that were significantly up- or down-regulated, respectively, in tumors from morphine-treated mice (*p*<0.05, fold change > 1.25). Many upregulated transcripts corresponded to structural ECM proteins previously associated with poor clinical outcomes in human PDAC and other solid cancers, including collagens 1, 3, and 5, as well as multiple laminin 5 subunits. Others were established ECM-modifying enzymes such as lysyl oxidase, metalloproteinases, cathepsins, and procollagen endopeptidases and proteinases **(Fig. 2A (red dots), Fig. 2B)**. To assess the most enriched pathways following morphine treatment, pathway analysis was performed using Reactome, GO, and KEGG databases. The top enriched pathways were predominantly associated with collagen formation and ECM organization, including “assembly of collagen fibrils,” “ECM organization,” and “collagen formation,” consistent with the increased collagen deposition and structural remodeling observed phenotypically **(Fig. 2C, D**, **Supp Fig. 2A)**. To assess the translational relevance of the morphine-induced upregulated genes, we developed a 23-gene “morphine-induced ECM signature” (MIE) derived from the morphine upregulated gene set **(Fig. 2E, see Methods for details on signature development)**. We next evaluated MIE signature scores across datasets representing well-established PDAC subtype classification systems. Elevated MIE signature scores were significantly enriched in aggressive PDAC subtypes, including the basal-like subtype (Moffitt), quasi-mesenchymal subtype (Collisson), and squamous subtype (Bailey), consistent with the poorly differentiated pathological assessment of tumors from morphine-treated mice [26–28] **(Fig. 2F)**. To determine the prognostic relevance of MIE, survival analyses were performed in PDAC cohorts. PDAC patients with elevated MIE scores exhibited significantly worse disease-free survival (DFS) and overall survival (OS) **(Fig. 2G)**. To further evaluate the broader clinical relevance of MIE, pan-cancer analyses were conducted across TCGA cohorts. Elevated MIE scores were associated with significantly worse DFS and OS across multiple cancer types, including uveal melanoma, mesothelioma, glioblastoma multiforme, low grade glioma, kidney renal papillary cell carcinoma, and cervical squamous cell carcinoma **(Fig. 2H, I, Supp Fig. 2B, C)**. Collectively, these results demonstrate that morphine treatment induces an ECM-related transcriptional program associated with more aggressive tumor phenotypes and poor clinical outcomes in PDAC and across multiple cancer types.

**Figure 2:**
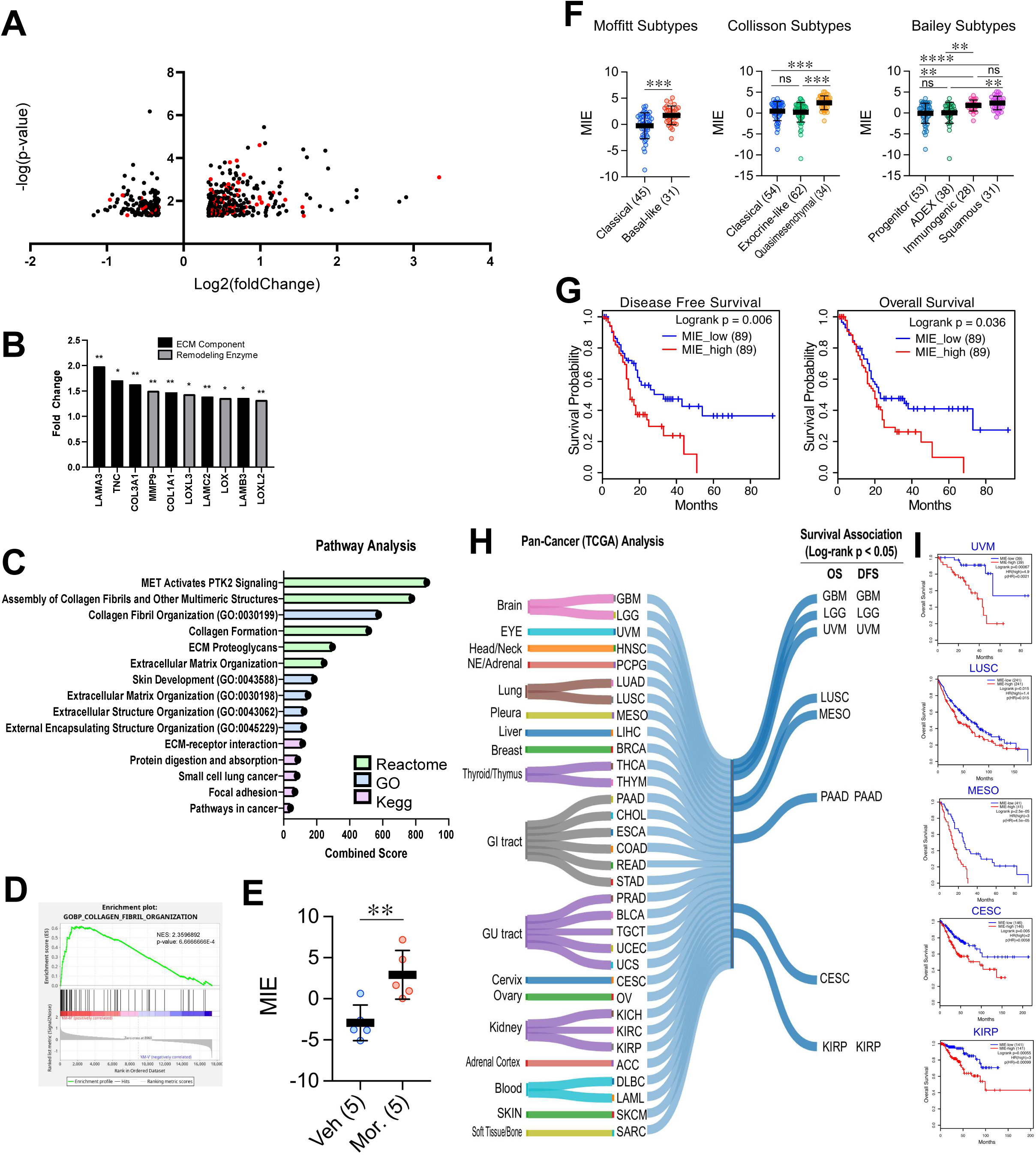
Morphine induces expression of an extracellular matrix signature associated with poor survival. **A,** Volcano plot of differentially expressed genes in tumors from morphine-treated mice relative to vehicle-treated mice with genes related to ECM organization highlighted in red. **B,** Upregulated extracellular matrix-related genes in tumors from morphine-treated mice relative to vehicle-treated controls. Statistical significance was determined based on differential expression analysis comparing tumors from morphine-treated and vehicle-treated mice. **C,** Enrichr combined scores of the top 5 enriched Reactome, GO, and KEGG terms in tumors from morphine-treated mice relative to vehicle-treated controls. **D,** Enrichment plot of GOBP_Collagen_Fibril_organization (*p* = 6.67E-4). **E,** MIE gene signature scores in tumors from morphine-treated vs vehicle-treated mice. An unpaired T test was used for analysis of statistical significance. **F,** MIE scores across PDAC subtypes defined by the Moffitt, Collisson, and Bailey classification systems. **G,** Elevated MIE predicts poor disease-free survival (DFS) and overall survival (OS) in TCGA-PAAD. **H,** Pan-cancer analysis of OS and DFS showing significant association between elevated MIE scores and poor clinical outcome (8 cancer types for OS, 5 cancer types for DFS). **I,** Representative Kaplan-Meier survival curves from cancer types shown in **H** demonstrating significant differences in OS based on MIE scores. Kaplan-Meier survival curves were used to estimate survival distributions, and statistical significance between groups was assessed using the log-rank (Mantel-Cox) test. Hazard ratios (HRs) were estimated using Cox proportional hazards regression models. *, *p* < 0.05; **, *p* < 0.01; ***, *p* < 0.001; **** *p* < 0.0001. Sample sizes are indicated in parenthesis.

### Opioid signaling regulates expression of ECM-related genes in CAFs in vitro

The high prevalence of ECM-related changes in our tumors from morphine-treated mice suggested that morphine may be acting directly on CAFs, the main producers of ECM in PDAC. To test this hypothesis directly, we sought to determine the role of opioid signaling in modulating the activity of pancreatic CAFs *in vitro*. First, we performed siRNA-mediated knockdown of the mu opioid receptor (OPRM1) in immortalized human pancreatic CAFs and observed significant decreases in the expression of the collagens COL1A1 and COL3A1 at the RNA and protein level **(Fig. 3A, B).** COL1A1 and COL3A1 were chosen for analysis because of their abundance and importance in the PDAC ECM, as well as their upregulation in tumors from morphine-treated mice **(Fig. 2B)** [8]. To replicate this pharmacologically, we treated CAFs with the OPRM1 antagonists naloxone and MNTX and observed significantly decreased expression of COL1A1 and COL3A1 at the RNA level **(Fig. 3C, Supp Fig. 3A)**. Interestingly, we also observed a reduction in expression of alpha smooth muscle actin (αSMA) following both OPRM1 knockdown as well as pharmacological OPRM1 inhibition **(Fig. 3A-C)**. Therefore, we provide evidence that opioid signaling directly regulates expression of critical PDAC CAF genes.

**Figure 3:**
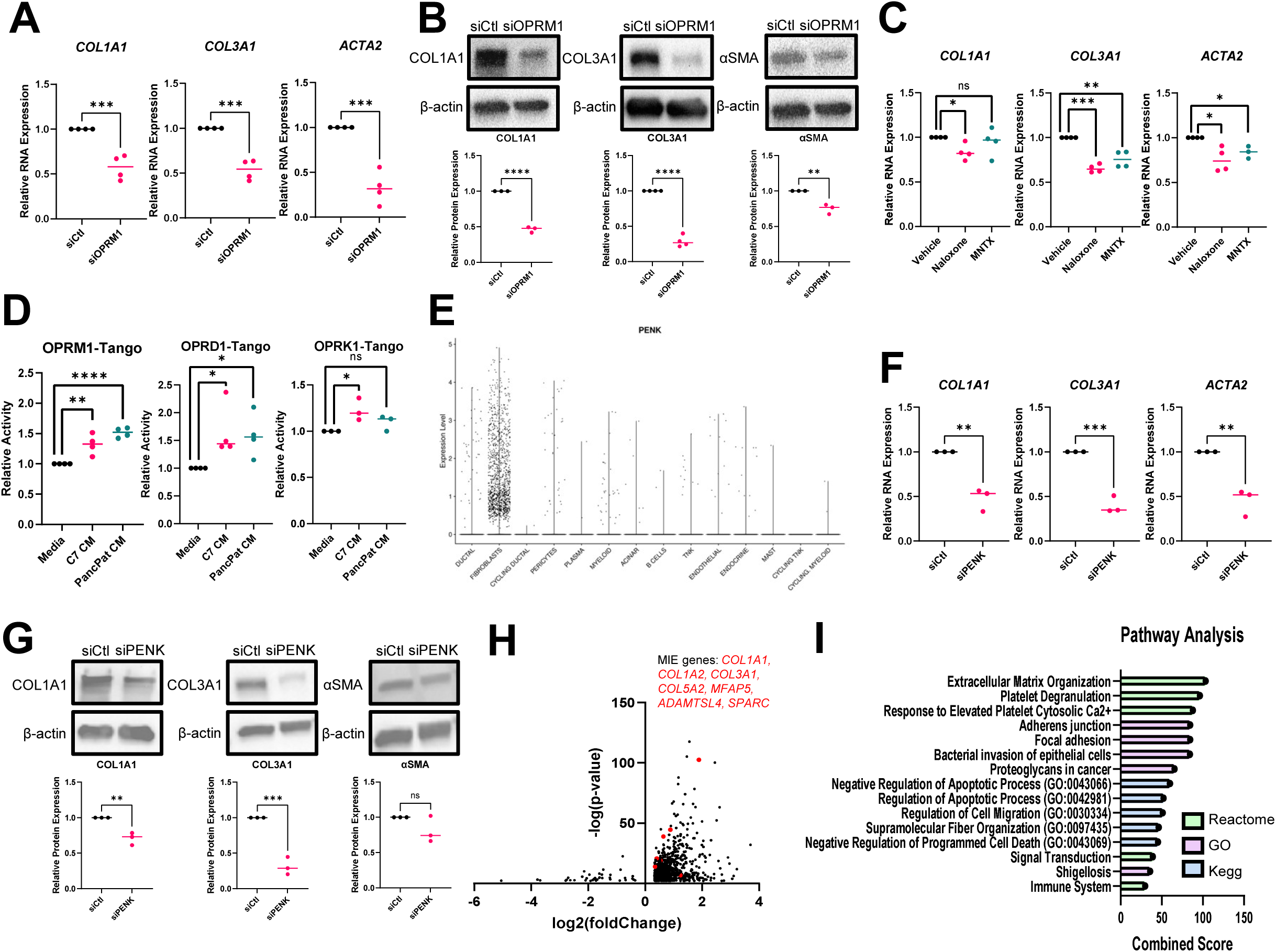
Endogenous opioids regulate expression of ECM-related genes in CAFs *in vitro*. **A,** qPCR for *COL1A1, COL3A1,* and *ACTA2* of siCtl or siOPRM1 immortalized human pancreatic C7-TA-PSC CAFs. **B,** Western blot of siCtl or siOPRM1 immortalized human C7-TA-PSC pancreatic CAFs for COL1A1, COL3A1, and αSMA (top) and respective quantifications (bottom). **C,** qPCR for *COL1A1, COL3A1,* and *ACTA2* of immortalized human pancreatic C7-TA-PSC CAFs treated with 20µM vehicle, naloxone, or MNTX for 6 hours. **D,** PRESTO-Tango Assays for OPRM1, OPRD1, and OPRK1 activation by conditioned media from immortalized human pancreatic CAFs. **E,** Expression of PENK by cell type in primary tumors using human single cell RNA sequencing data. **F,** qPCR for *COL1A1, COL3A1,* and *ACTA2* of siCtl or siPENK immortalized human pancreatic CAFs. **G,** Western blot of siCtl or siPENK immortalized human pancreatic CAFs for COL1A1, COL3A1, and αSMA (top) and respective quantifications (bottom). **H,** Volcano plot of upregulated and downregulated genes in PENK^+^ CAFs from human single cell RNA sequencing data. MIE signature genes are highlighted in red. **I,** Enrichr combined scores of the top 5 enriched Reactome, GO, and KEGG terms from upregulated genes in PENK^+^ CAFs. Unpaired T tests were performed for the comparison of two groups for statistical significance. *, *p* < 0.05; **, *p* < 0.01; ***, *p* < 0.001; **** *p* < 0.0001.

### PDAC CAFs secrete endogenous opioids regulating expression of ECM genes

As OPRM1 inhibition decreased collagen and αSMA expression in the absence of an exogenous agonist, we hypothesized that CAFs produce endogenous opioid peptides capable of driving collagen expression. To test this, we utilized the PRESTO-Tango assay, a luciferase-based method of measuring G-protein coupled receptor activation [29]. Conditioned media (CM) from two immortalized human pancreatic CAF lines significantly increased activation of both OPRM1 and OPRD1, but not OPRK1 **(Fig. 3D)**. To investigate which endogenous opioids could be produced by CAFs, we analyzed single cell RNA sequencing data from human pancreatic tumors [30] for expression of the three endogenous opioid peptide precursors: POMC, which produces the endogenous opioid peptide beta-endorphin; PDYN, which produces dynorphin; and PENK, which produces enkephalin peptides [31]. We found that PENK was predominantly expressed in fibroblasts relative to other cell types in primary tumors, with increased expression compared to adjacent normal **(Fig. 3E, Supp Fig. 3B)**. In contrast, neither POMC nor PDYN were as highly expressed in fibroblasts relative to other cell types, nor showed significant changes in expression between normal and tumor **(Supp Fig. 3C-F)**. To determine the functional relevance of PENK in CAFs, we knocked down PENK expression by siRNA in immortalized pancreatic CAFs and observed a significant reduction in COL1A1, COL3A1, and αSMA expression at both the RNA and protein level **(Fig. 3F, G, Supp Fig. 3G)**. To further elucidate the role of PENK in CAFs, we again analyzed human single cell RNA sequencing data to determine differentially expressed genes based on PENK expression. We found 1097 upregulated genes and 41 downregulated genes in PENK^+^ CAFs. Intriguingly, expression of 7 of the 23 genes present in our MIE gene signature positively correlate with high expression of PENK, as highlighted in red **(Fig. 3H)**. Pathway analysis of the upregulated genes in PENK^+^ CAFs revealed Extracellular Matrix Organization as the top enriched pathway **(Fig. 3I)**, consistent with the high prevalence of ECM-related pathways in the RNA sequencing results from tumors from morphine-treated mice in our *in vivo* study **(Fig. 2C)**. Taken together, our results demonstrate that PDAC CAFs secrete endogenous opioids, primarily driven by expression of the opioid precursor PENK, capable of driving collagen expression in an autocrine manner.

### Methylnaltrexone reduces tumor weight, aggressiveness, and opioid-induced ECM remodeling

The reduction in collagen expression following MNTX treatment of CAFs *in vitro* suggested its potential applicability to target endogenous and exogenous opioid-induced ECM remodeling *in vivo*. Importantly, as MNTX does not cross the blood-brain barrier, it is used clinically in patients concurrently with opioids to treat opioid-induced constipation. To test this directly, we orthotopically implanted KPC tumor pieces into the pancreata of wild-type C57BL/6 mice, allowed the tumors to establish for three days, then treated the mice with morphine alone, MNTX alone, morphine and MNTX in combination, or vehicle control for 12 days **(Fig. 4A)**. Histopathological analysis revealed notable differences in both tumor epithelium and stroma between the four groups. As observed in our earlier experiment **(Fig. 1B, C)**, tumors from vehicle-treated mice were predominantly well-to-moderately differentiated, whereas tumors from morphine-treated mice were dominantly moderate-to-poorly differentiated. Tumors from mice treated with MNTX and the combination also displayed increased moderate-to-poor differentiation when compared to vehicle **(Fig. 4B, Supp Fig. 4A-C)**. Tumors from morphine-treated mice exhibited increased levels of typical and atypical mitoses, characteristic of more aggressive disease. Strikingly, the combination of MNTX and morphine treatment was able to restore levels of mitoses to those observed in tumors from vehicle-treated mice **(Fig. 4B)**. We next assessed collagen deposition by performing Masson’s trichrome and picrosirius red staining. As in our earlier experiment, **(Fig. 1F)**, increased collagen was observed in the tumors from morphine-treated mice. In contrast, tumors from the MNTX- and combination-treated mice displayed reduced staining for collagen **(Fig. 4C)**. To assess changes in the polymerization status of collagen fibers between the groups, we used multiphoton microscopy for SHG and found a statistically significant increase in collagen fiber density in the tumors from morphine-treated mice **(Fig. 4D)**, indicating increased collagen bundling and maturation, consistent with our previous results **(Fig. 1G)**. There was also a statistically significant increase in collagen fiber density in tumors from MNTX-treated mice. Notably, tumors from combination-treated mice exhibited similar levels of collagen fiber density as seen in tumors from vehicle-treated mice **(Fig. 4D)**. Alterations in collagen bundling and maturation are associated with differences in PDAC aggressiveness and disease outcomes. Accordingly, tumors from combination-treated mice weighed significantly less than control tumors or tumors from mice treated with morphine alone **(Fig. 4E)**. Most strikingly, MNTX, either alone or in combination with morphine, decreased the presence of ascites, a feature associated with more aggressive disease **(Fig. 4F)**. Collectively, these results demonstrate that MNTX, both alone and in combination with morphine, alters the PDAC tumor microenvironment in a manner associated with reduced PDAC aggressiveness. We propose a model whereby both exogenous and endogenous opioids act on CAF-resident opioid receptors to drive tumor-promoting ECM remodeling, associated with poor patient outcome **(Fig. 4G)**. Furthermore, we propose that the FDA-approved peripherally restricted OPRM1 antagonist MNTX represents a novel approach to counteract these effects and reduce tumor aggressiveness.

**Figure 4:**
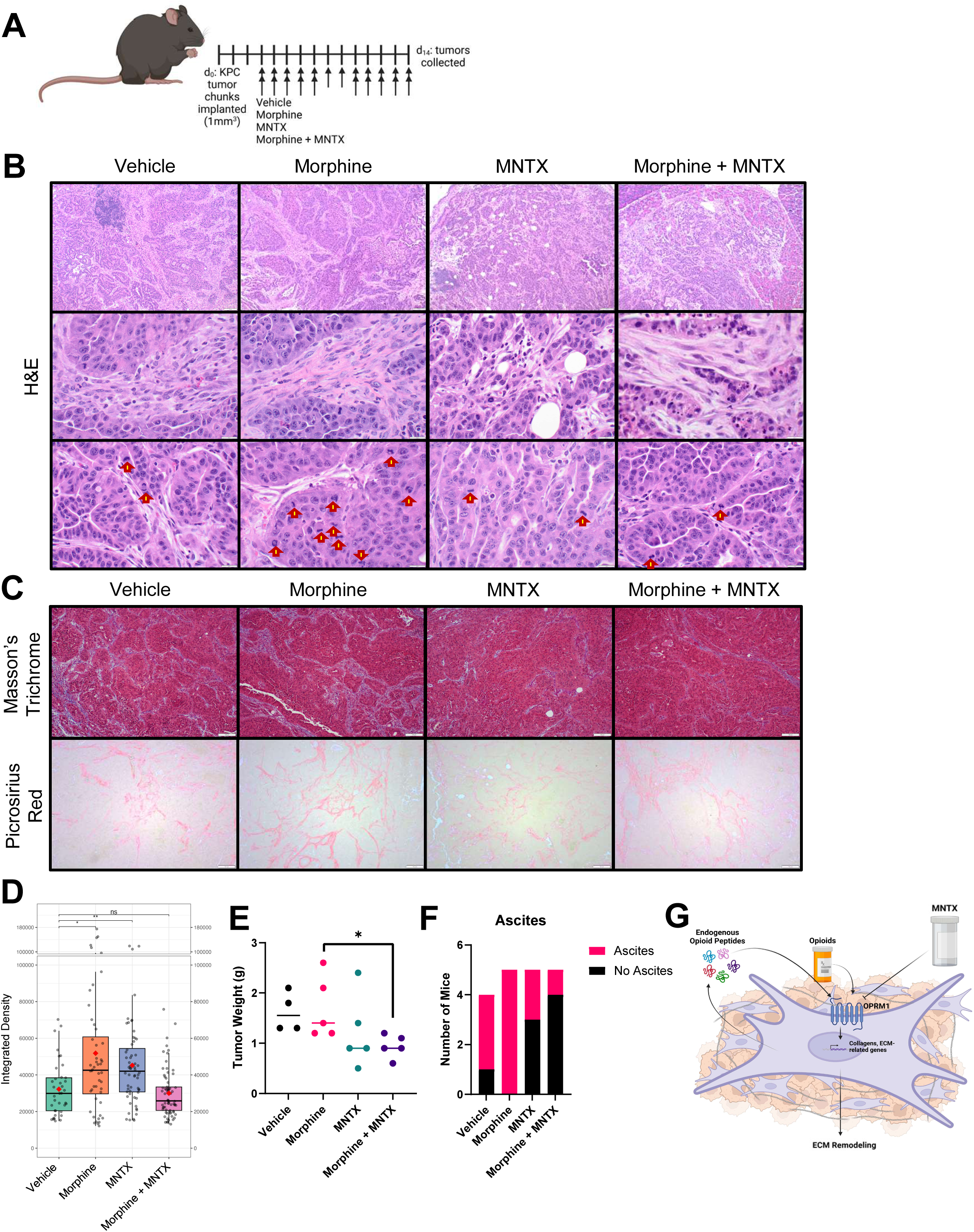
MNTX reduces tumor weight, aggressiveness, and ECM remodeling driven by both exogenous and endogenous opioids. **A,** Experimental schematic of orthotopic transplantation of LSL-KrasG12D/+; LSL-Trp53R172H/+; Pdx-1-Cre (KPC) tumors into C57BL/6 mice and treatment with subcutaneous vehicle (*n* = 4), morphine (10mg/kg) (*n* = 5), MNTX (1mg/kg) (*n* = 5), or morphine + MNTX (10mg/kg and 1mg/kg) (*n* = 5) for 12 days. Arrows indicate doses on each individual day. **B,** Representative 10x (top) and 60x (middle and bottom) H&E images of tumors from (left to right) vehicle, morphine, MNTX, and morphine + MNTX-treated mice. Bottom images demonstrate typical and atypical mitoses using yellow/red arrows. Scale bars represent 100µm (top) and 20µm (middle and bottom). **C,** Representative 10x images of Masson’s Trichrome (top) and Picrosirius Red (bottom) staining of tumors from (left to right) vehicle, morphine, MNTX, and morphine + MNTX-treated mice. Scale bars represent 100µm. **D,** Quantification of collagen fiber density by SHG microscopy in tumors from vehicle, morphine, MNTX, and morphine + MNTX-treated mice. Statistical comparisons were made using the Wilcoxon test. **E,** Weight of tumors (g) from mice treated with vehicle, morphine, MNTX, and morphine + MNTX. Unpaired T tests were used for the comparison of two groups for analysis of statistical significance. **F,** Ascites burden in vehicle, morphine, MNTX, and morphine + MNTX treated mice. **G,** Graphical model. *, *p* < 0.05; **, *p* < 0.01; ***, *p* < 0.001; **** *p* < 0.0001.

## Discussion

Patients with cancer receive a plethora of medications to combat both tumor-intrinsic and treatment-induced side effects. However, the impact of these medications on tumor progression and treatment response is frequently overlooked [32]. Here, we demonstrate that the commonly prescribed opioid, morphine, has profound impacts on pancreatic tumor biology, and in particular, regulation of the ECM. Specifically, we provide the first evidence that morphine drives ECM remodeling in the PDAC TME **(Fig. 1 and 4)** and promotes expression of an ECM gene signature that correlates with worse prognosis in pancreatic cancer and other tumor types **(Fig. 2)**. We find that morphine treatment promotes more poorly differentiated tumors, with increased collagen bundling and maturation, features associated with poor prognosis **(Fig. 1)**. Importantly, we demonstrate that treatment with the FDA-approved peripherally restricted OPRM1 antagonist MNTX reverses opioid-induced ECM alterations and reduces tumor aggressiveness **(Fig. 4)**. We also provide the first evidence that CAFs produce endogenous opioids, driving expression of collagens and ECM remodeling enzymes **(Fig. 3)**. In summary, we propose that both endogenous and exogenous opioids promote tumor aggressiveness and ECM remodeling, and suggest that MNTX, an FDA-approved drug already in use in PDAC patients, may provide therapeutic benefit.

Patients with cancer are prescribed a variety of medications while undergoing treatment for their disease, including medications prescribed both before and after diagnosis. Many of these commonly prescribed therapeutics target G-protein coupled receptors (GPCRs), which are critical regulators of tumor biology [33–35]. Thus, there is growing interest in investigating the impact of medications taken by patients with cancer on disease progression and therapy response [36]. For example, β-adrenergic signaling promotes tumor progression through diverse processes, including promoting proliferation, angiogenesis, invasiveness, impaired immune function, and metastasis in various cancer types. Accordingly, β-blockers have shown promise in inhibiting these processes and improving outcomes [37–41]. Further, recent work has elucidated the role of the benzodiazepine lorazepam in promoting pancreatic tumor aggressiveness and correlating with worse patient outcomes, whereas alprazolam correlates with improved progression-free survival and features of less aggressive disease [42, 43]. Thus, it is well established that GPCR-targeted drugs can have profound effects outside of their intended functions, with important implications for patient outcome [32]. Our study contributes to the current body of literature investigating the impact of commonly prescribed medications by identifying a previously unknown tumor-promoting impact of opioids on the pancreatic cancer phenotype.

Opioids are the most commonly prescribed medications for managing moderate-to-severe cancer-related pain [44]. In pancreatic cancer specifically, approximately 75% of patients are prescribed opioids regardless of disease stage [19]. Opioid analgesics relieve pain through activating opioid receptors, most commonly the GPCR mu opioid receptor (OPRM1), resulting in cellular hyperpolarization and inhibition of pain transmission [45, 46]. Both the use of opioids as well as the increased expression of OPRM1 is associated with worse prognosis across various cancer types, including pancreatic cancer [20, 21]. Previous studies have investigated the direct effects of opioids on tumor cells. Specifically, OPRM1 regulates epithelial-mesenchymal transition (EMT), migration and invasion, tumor cell proliferation, and angiogenesis [47–49]. Opioids also increase stemness of pancreatic tumor cells, as well as promote resistance to gemcitabine and 5-fluorouracil [22]. Opioids may also promote tumor progression through direct effects on the immune component of the TME. Opioid use correlates with poor response to immunotherapy in multiple tumor types [50–53]. McIlvried et al. demonstrated a potential mechanism for this through OPRM1-mediated suppression of CD8+ T cells, resulting in impaired response to anti-PD-1 checkpoint inhibition in head and neck cancer [24]. While studies have interrogated the direct effects of opioids on PDAC tumor cells *in vitro*, studies utilizing *in vivo* models are limited. One recent study investigating the effects of morphine on PDAC involved the use of a subcutaneous model system that lacks the relevant ECM [54]. Additionally, anti-tumorigenic effects of tramadol, a weak OPRM1 agonist, have been reported. However, tramadol also acts as an inhibitor of monoamine neurotransmitter reuptake, indicating that the observed effects, specifically those unaltered in the presence of the OPRM1 antagonist naltrexone, may not be specific to opioid signaling [55–57]. Although our study focuses on the role of opioids in CAFs, direct and indirect effects on tumors cells and immune cells cannot be ruled out. Our study is the first to link opioid signaling to regulation of the tumor ECM and CAF biology, critical aspects of pancreatic cancer progression and therapy response. Therefore, we provide novel insight into how a commonly prescribed drug impacts tumor biology and uncover a novel endogenous pathway by which CAFs regulate the ECM.

Outside the context of cancer, the effect of opioids on fibrosis and the ECM is well-appreciated. In opioid use disorder, opioids remodel the ECM in the brain through increased expression of ECM remodeling enzymes MMP-2 and MMP-9 [58]. Opioids also increase the collagen content of wounds in mice [59]. Similarly, opioid treatment increases activation of fibroblasts and persistence of myofibroblasts at the end of wound healing [60]. Morphine promotes cardiac interstitial fibrosis in mice, which is associated with cardiac damage and remodeling [61]. Further, morphine induces the accumulation of collagens in renal interstitial cells *in vitro*, which may contribute to renal injury and interstitial scarring [62]. Intriguingly, opioids exacerbated a mouse model of pancreatitis, as evidenced by increased fibrosis in addition to increased necrosis and infiltration of immune cells [63]. This is especially relevant, as chronic pancreatitis is a known risk factor for the development of pancreatic cancer. Endogenous opioid peptides have also been implicated in driving fibrosis, outside of the context of cancer. For example, the OPRM1 antagonist naloxone reduced liver fibrosis in a dimethylnitrosamine (DMN)-induced rat model of hepatic fibrosis [64]. This study did not involve the addition of an exogenous agonist, indicating that the reduced fibrotic effects observed are due to the inhibition of endogenously produced opioid peptides. Endogenous opioid peptides are produced by the cleavage of three main precursors, proenkephalin (PENK) which produces enkephalin peptides, proopiomelanocortin (POMC) which produces beta-endorphin, and prodynorphin (PDYN) which produces dynorphins. These peptides signal through three GPCR opioid receptors, OPRM1, the delta opioid receptor (OPRD1), and the kappa opioid receptor (OPRK1), with enkephalins having the strongest affinity for OPRD1 and OPRM1, beta-endorphin having the strongest affinity for OPRD1 and OPRM1, and dynorphins having the strongest affinity for OPRK1 [31]. Previous studies on the role of endogenous opioid peptides in cancer are limited. Several studies suggest an anti-tumorigenic role for enkephalins, the cleavage product of PENK; however, the mechanism by which this occurs involves the zeta opioid receptor, also known as the opioid growth factor receptor, which is structurally distinct from the classical opioid receptors. Further, currently published studies focus exclusively on the role of PENK in tumor cells and immune cells, whereas our study is the first to focus on CAFs [65–70]. Although our study focuses solely on the role of PENK in CAFs, we acknowledge the possibility that CAFs, among other cell types in the pancreatic TME, also exhibit expression of POMC, which could be driving similar effects. Additionally, the existence of a fifth opioid receptor, the nociceptin receptor (OPRL1), has been of recent interest. However, this receptor is not sensitive to prescription opioids or their antagonists, and is therefore outside the scope of the current study [71]. While both exogenous and endogenous opioids have been implicated in promoting fibrosis and collagen deposition in normal fibroblasts and in noncancerous physiological settings, there are no studies addressing this in the context of cancer. To our knowledge, we are the first study to investigate this phenomenon in the context of cancer biology and to uncover a mechanism for CAFs in driving both exogenous and endogenous opioid-induced tumor aggressiveness.

Targeting CAFs and the ECM has been of growing interest in the pancreatic cancer field due to the immense influence of the stroma on tumor progression and therapeutic resistance. Following the discovery that depleting the stroma can actually promote tumor aggressiveness, recent work has focused on stromal normalization, rather than depletion, as a goal to improve therapeutic efficacy [72, 73]. For example, agonism of the vitamin D receptor results in reprogramming of CAFs to a more quiescent state, leading to stromal remodeling, reduced tumor volume, and increased survival in mice [74]. Further, the anti-microtubule agent eribulin can induce CAF normalization, resulting in reduced pancreatic tumor cell invasion and viability *in vitro* [7]. Directly targeting the structure of the ECM can improve drug delivery and therapeutic responses. Depletion of the tumor-promoting ECM component hyaluronic acid resulted in improved therapeutic efficacy of gemcitabine in mice [75]. Further, studies have also investigated methods of targeting the remodeling of the ECM. For example, inhibition of the lysyl oxidase family of enzymes involved in collagen crosslinking reduces cancer cell invasion and improves survival in combination with gemcitabine in KPC mice [76]. Strikingly, treatment with the commonly prescribed angiotensin II receptor blocker Losartan reduces collagen I and hyaluronic acid production and leads to improved chemotherapeutic efficacy and reduced tumor volume in an orthotopic mouse model of pancreatic cancer [77]. CAFs and the ECM have complex roles in promoting tumor progression and therapeutic resistance. Methods of targeting these processes, either through CAF normalization, degrading ECM components, or inhibiting ECM remodeling have shown promise in improving therapeutic efficacy in preclinical models [8, 78–80]. Our study is the first to both elucidate a role for opioids in driving CAF-induced ECM remodeling as well as a method to target and reduce opioid-induced ECM alterations with an FDA-approved therapeutic, already in use in PDAC patients.

There are currently three FDA-approved peripherally restricted OPRM1 antagonists (PAMORAs): methylnaltrexone (MNTX), naldemedine, and naloxegol, so named for their inability to cross the blood-brain barrier. MNTX, the most extensively studied of the three in the current body of literature, is a methylated derivative of naltrexone, an OPRM1 antagonist that readily crosses the blood-brain barrier [81, 82]. MNTX was approved by the FDA for the treatment of opioid-induced side effects such as constipation and is the only PAMORA without clinically relevant cytochrome P450 interactions [81, 83]. Despite the growing body of literature suggesting a role for opioids and OPRM1 signaling in promoting cancer progression, there has been limited research into the use of PAMORAs as potential therapeutics in cancer [84]. *In vitro* studies demonstrate that MNTX can have antiproliferative effects directly on tumor cells [85], and MNTX reduced tumor growth in a mouse model of lung carcinoma [23]. Co-administration of MNTX inhibited the morphine-induced immunosuppression of CD8+ T cells [24]. Importantly, a retrospective analysis of patients with advanced cancer, all of whom were prescribed opioids, found that those who had received MNTX had significantly improved overall survival compared to those who received the placebo [86]. To our knowledge, the present study is the first to demonstrate MNTX as a therapeutic for the inhibition of both exogenous and endogenous opioid-induced alterations in the ECM and tumor aggressiveness in pancreatic cancer.

In summary, we have investigated the effect of opioids on the PDAC TME. We have discovered that opioids drive ECM remodeling and more poorly differentiated tumors, indicative of increased tumor aggressiveness. Similarly, we uncovered the ability of pancreatic CAFs to produce endogenous opioids as well as their role in ECM remodeling. Importantly, we propose a method for targeting this exogenous and endogenous opioid-induced tumor aggressiveness by repurposing an FDA-approved peripherally restricted OPRM1 antagonist, MNTX. Because of the high frequency of opioid prescriptions in pancreatic cancer patients regardless of stage, our findings are broadly relevant. Importantly, there is the potential for rapid translational applicability to the clinic, as MNTX is already safely used in pancreatic cancer patients for the treatment of opioid-induced side effects. Lastly, we believe this study could be applicable to other tumor types that are characterized by the prevalence of CAFs or a dense, fibrotic ECM.

## Authors’ Disclosures

EC and JFB collaborate with Visiopharm and Lunaphore/BioTechne on projects unrelated to this study.

## Authors’ Contributions

**K.E. Maraszek:** Conceptualization, investigation, writing-original draft, writing-review and editing.

**A.A. Tisdale:** Investigation. **H.D. Reavis**: Investigation. **X. Liu**: Investigation, formal analysis, methodology. **E. Cortes Gomez**: Formal analysis. **A. Dolskii**: Investigation, formal analysis, methodology. **C. Brown**: Investigation, formal analysis. **D. Thakkar**: Investigation. **M. Dungan**: Investigation. **A. Adhikari**: Investigation. **E. Mackey**: Formal analysis. **J. Franco-Barraza**: Formal analysis, supervision, methodology. **B.A. Pereira:** Formal analysis. **P. Timpson:** Supervision.

**D.G. Tang**: Supervision. **N. Steele**: Formal analysis. **E. Cukierman**: Formal analysis, resources, supervision, editing. **M.E. Feigin**: Resources, formal analysis, supervision, funding acquisition, project administration, writing-review and editing.

## Acknowledgements

We would like to thank Dr. Brian Roth for the generous donation of cell lines that were used in this study. We would also like to thank Amanda Tracz for her assistance with the orthotopic KPC mouse model used in our *in vivo* studies. Research in this study through the Roswell Park Comprehensive Cancer Center’s Comparative Oncology Shared Resource, Biorepository and Laboratory Services Shared Resource, Experimental Tumor Models Shared Resource, Genomics Shared Resource, and Bioinformatics Shared Resource was supported through the NCI grant P30CA016056. This study was also supported by funding from the ELF Foundation to MEF. We dedicate this work to Patricia Keely (a pioneer in ECM biology). Added support was from the 5th AHEPA Family Cancer Research Foundation, the Pancreatic Cancer Cure Foundation, Mrs. Marina P. Zazanis and the Zazanis family, the M&C Greenberg Pancreatic Cancer Institute to EC. Grants: R01CA269660, U54CA272686, S10ODO23666, DOD HT9425-23-1-0584, the ACS Wilmott Family Pancreatic Cancer Professorship RP-23-1070169-01-WRP, and the NIH/NCI Comprehensive Cancer Center Core Grant P30CA06927, supporting the Cell Culture, Imaging, and Histopathology facilities.

## Materials and Methods

### Murine experiments

Mice were housed and maintained in the Comparative Oncology Shared Resource at Roswell Park Comprehensive Cancer Center. All experiments were conducted under IACUC protocol #1381M.

### LSL-KrasG12D/+; LSL-trp53r172h/+; pdx-1-cre (KPC) orthotopic allograft short-term morphine study

A KPC002 allograft derived from a female KPC mouse was stored in freezing media (50% RPMI, 40% FBS, and 10% DMSO) in liquid nitrogen. The allograft tissue was then passaged once in strain-matched C57BL/6 female mice by dipping a 1-2mm^3^ piece of tumor tissue in Matrigel Matrix Basement Membrane (Corning, cat. #354234) and implanting subcutaneously. The tumor tissue was harvested about a week and a half later. ∼1mm^3^ tumor pieces were implanted orthotopically into 20 C57BL/6 female mice. Mice received carprofen as an analgesic in place of buprenorphine. Mice were given a single carprofen injection (5mg/kg, subcutaneous) at the time of surgery. The next day, a carprofen tablet (Bio-Serv, cat. #SMD150-2) was placed in each cage. Three days post-implantation, each mouse was treated with either 10mg/kg morphine (Patterson Veterinary, cat. #07-892-4699) or vehicle control (n = 10 per arm) twice daily on weekdays and once daily on weekends by subcutaneous (s.c.) injection. Morphine was prepared by dilution in sterile saline. Mice were sacrificed on day 14 of treatment.

### LSL-KrasG12D/+; LSL-trp53r172h/+; pdx-1-cre (KPC) orthotopic allograft short-term MNTX study

A KPC002 allograft derived from a female KPC mouse was stored in freezing media (50% RPMI, 40% FBS, and 10% DMSO) in liquid nitrogen. The allograft tissue was then passaged in strain-matched C57BL/6 female mice as described in the section above. ∼1mm^3^ tumor pieces were implanted orthotopically into 25 C57BL/6 female mice. Mice received carprofen as an analgesic in place of buprenorphine. Three days post-implantation, each mouse was treated with either 10mg/kg morphine, 1mg/kg methylnaltrexone bromide (MNTX) (Cayman Chemical, cat. #24072), 10mg/kg morphine and 1mg/kg MNTX combined, or vehicle control (n = 5 per arm). Morphine was prepared as described in the previous section. MNTX was prepared fresh daily by diluting 1mg vials in 1mL of PBS to make a 1mg/mL stock solution, which was then further diluted with PBS to 0.1mg/mL. Morphine was administered twice daily on weekdays and once daily on weekends, while MNTX was administered once daily. Mice were sacrificed on day 12 of treatment.

### H&E staining

Freshly isolated tumors were fixed in 10% neutral buffered formalin solution (Sigma-Aldrich, cat. #HT501128) for 24 hours prior to processing. Tumor processing and H&E staining was performed in the Biorepository and Laboratory Services (BLS) Shared Resource at Roswell Park. Tumor processing was performed using a HistoCore Arcadia (H+C) (Leica Biosystems) embedder and sliced in 4µm sections using an Olympus Cut 4060 Microtome. FFPE unstained slides were rehydrated as follows: xylene: 5 minutes (repeat 3 times), 100% ethanol: 1 minute (repeat 2 times), 95% ethanol: 45 seconds, 70% ethanol: 30 seconds. Slides were then rinsed in running tap water across multiple wash stations (2-4 minutes total) to ensure complete removal of alcohol. Slides were stained using an automated H&E stainer (Leica Autostainer XL). Nuclear staining was performed with hematoxylin for 3 minutes, followed by differentiation in acid alcohol for 1 minute and subsequent bluing in a bluing solution for 45 seconds. Cytoplasmic counterstaining using eosin was performed for 30 seconds. Slides were then dehydrated using 95% ethanol, followed by multiple changes of 100% ethanol. Ethanol was cleared using 3 changes of Histoclear (2 minutes each). Slides were then prepared for mounting and cover slipped.

### Masson’s trichrome staining

Freshly isolated tumors were fixed in 10% neutral buffered formalin solution (Sigma-Aldrich, cat. #HT501128) for 24 hours prior to processing. Tumor processing was performed in the Biorepository and Laboratory Services (BLS) Shared Resource at Roswell Park as described above. FFPE slides were rehydrated as follows: xylene: 3 minutes (repeat 3 times), 100% ethanol: 3 minutes (repeat 3 times), 95% ethanol: 3 minutes, 70% ethanol: 3 minutes, distilled water: 5 minutes. The Abcam trichrome stain kit (Abcam, cat. #ab150686) was then used according to the manufacturer’s instructions. For step 9, the slides were washed with distilled water for 2 minutes. In Step 12, the slides were washed with distilled water for 30 seconds. The slides were dehydrated as follows: 95% ethanol: 3 minutes (repeat 2 times), 100% ethanol: 3 minutes (repeat 2 times), and xylene: 5 minutes (repeat 3 times). The slides were left in xylene overnight, dried briefly, and cover-slipped using Poly-Mount.

### Picrosirius red staining

Freshly isolated tumors were fixed in 10% neutral buffered formalin solution (Sigma-Aldrich, cat. #HT501128) for 24 hours prior to processing. Tumor processing was performed in the Biorepository and Laboratory Services (BLS) Shared Resource at Roswell Park as described above. FFPE slides were rehydrated as follows: xylene: 3 minutes (repeat 3 times), 100% ethanol: 3 minutes (repeat 3 times), 95% ethanol: 3 minutes, 70% ethanol: 3 minutes, distilled water: 5 minutes. The Abcam Picro Sirius red stain kit (Abcam, cat. #ab150681) was then used according to the manufacturer’s instructions. In step 3, slides were rinsed in each change of acetic acid solution for 2 minutes. In step 5, slides were dehydrated in each change of absolute alcohol for 3 minutes. Slides were then placed in three changes of xylene for 3 minutes each. Slides were dried briefly, and cover-slipped using Poly-Mount.

### Immunohistochemistry

Freshly isolated tumors were fixed in 10% neutral buffered formalin solution (Sigma-Aldrich, cat. #HT501128) for 24 hours prior to processing. FFPE slides were rehydrated as follows: xylene: 5 minutes (repeat 3 times), 100% ethanol: 10 minutes, 95% ethanol: 10 minutes (repeat 2 times), 70% ethanol: 10 minutes, distilled water: 5 minutes (repeat 2 times). For antigen retrieval, rehydrated slides were placed in slide chambers with pH 6.0 1x Antigen Unmasking Solution, Citric Acid Based (Vector Laboratories, cat. #H-3300) in the 2100 Retriever (Aptum Biologics Ltd) and the antigen-unmasking cycle was run according to the manufacturer’s instructions. Following completion of the unmasking cycle, slides were allowed to cool overnight. Slides were washed 3x with distilled water and incubated in 1x PBS for 15 minutes. Endogenous peroxidase activity was blocked for 10 minutes using 3% lab-grade hydrogen peroxide (Fisher Science Education, cat. #S25359). Slides were washed using 1x PBS (0.1% Tween) and then blocked with normal goat serum (2.5%) for 20 minutes. For primary antibody incubation, slides were incubated overnight at 4°C. Primary antibodies were diluted in normal goat serum (2.5%) from the ImmPRESS HRP Goat Anti-Rabbit IgG Polymer Detection Kit, Peroxidase (Vector Laboratories, cat. #MP-7451-15). Primary antibodies include the following: Ki67 rabbit polyclonal antibody (Novus Biologicals, cat. #NB600-1209, 1:300 dilution) and αSMA rabbit polyclonal antibody (Proteintech, cat. #14395-1-AP, 1:1500 dilution). The following day, slides were washed with PBS (0.1% Tween) for 5 minutes. Slides were then incubated with ImmPRESS Polymer reagent for 30 minutes. Following this, slides were washed twice with PBS (0.1% Tween) for five minutes and then incubated for 30 seconds with the DAB Substrate Kit, Peroxidase (HRP; Vector Laboratories, cat. #SK-4100) according to the manufacturer’s instructions. After development, slides were counterstained with hematoxylin. Slides were dehydrated as follows: 95% ethanol: 3 minutes, 100% ethanol: 3 minutes, xylene: 3 minutes (repeat 3 times). Slides were dried briefly, and cover-slipped using Poly-Mount.

### Histopathological evaluation of differentiation status

Histopathological analysis was conducted by visually assessing tumor areas and the degree of expression of features. To support this evaluation and enable structural assessment of tissue architecture, all H&E-stained specimens were digitized using a NanoZoomer S60 high-throughput slide scanner (Hamamatsu Photonics, Japan). Sections were scanned under brightfield illumination to generate high-resolution whole-slide images. A blinded histopathologic evaluation was done for each individual sample of the different groups. The grade of the tumor (based on the extent of glandular differentiation) was noted indicating whether they were well-differentiated (>95% tumor composed of glands), moderately differentiated (50-95% glands) or poorly differentiated (< 50% glands) according to the TNM histologic grading system in the Protocol for the Examination of Specimens from Patients with Carcinoma of the Pancreas (College of American Pathologists). The surrounding stroma was also evaluated for the presence and extent of fibrosis as well as the presence or absence of inflammation. The findings are represented as a percentage of the tissue sample that was evaluated.

### Second Harmonic Generation (SHG) microscopy

Mature collagen fibers in H&E-stained mouse pancreatic sections were analyzed using Second Harmonic Generation (SHG) microscopy. Imaging was performed on a Leica TCS SP8 DIVE Multiphoton System (Leica Microsystems, Germany) equipped with an HC Fluotar L25x/0.95 W water immersion objective. The collagen fibers were excited using an IR pulsing laser Chameleon (Coherent Inc.) set to 860 nm. Backward-scattered SHG signal was collected with a high-sensitivity HyD NDD detector within a spectral range of 420–440 nm. Under the supervision of a trained pathologist, two regions of interest (ROIs) per specimen were selected to ensure representative tissue sampling, with each ROI consisting of 4–6 tiles. SHG image acquisition was performed using the automated function of Leica Application Suite X (LAS X) software (v3.5.5) with standardized settings applied uniformly across all ROIs. Collagen-specific SHG signatures were captured as monochromatic 16-bit image stacks spanning the entire tissue thickness with a Z-step size of 1 µm.

### SHG image analysis

For each histological slide, two representative regions of interest (ROIs) were manually selected at the interface between neoplastic tissue and normal regions, based on visual evidence of increased collagen deposition (desmoplasia) and verified by a pathologist. Each ROI provided a comprehensive representation of the extracellular matrix architecture. Brightfield microscopy was performed alongside SHG imaging to ensure anatomical accuracy and enable precise correlation of collagen fiber distribution with underlying histopathological features of the pancreatic tissue. To ensure reproducibility and systematic data handling, all raw microscopy images in Leica Image File (.lif) format were processed using an automated Python-based pipeline. The workflow used the *readlif* library for metadata extraction and PyImageJ to interface with a headless instance of ImageJ2/Fiji [ImageJ: https://imagej.net/software/imagej/ [87]; FIJI: https://fiji.sc [88]]. During processing, each tile was saved as a standardized 16-bit TIFF file. After data conversion, quantitative analysis of signal intensity and area for SHG collagen bundles was performed. For each dataset, a Maximum Intensity Projection (MIP) was generated to consolidate 3D collagen structures into a single 2D plane, standardized to a resolution of 1024 x 1024 pixels. SHG-positive collagen fibers were segmented using a thresholding algorithm with a consistent fixed manual range to generate binary masks. These masks were converted to 16-bit depth and applied to the original MIP images using a bitwise AND operation, ensuring intensity measurements were derived exclusively from pixels within the segmented collagen bundles while preserving original bit-depth values (0-65535 grayscale levels). Particle area was quantified in calibrated spatial units based on image metadata, and integrated density (IntDen) was calculated as the sum of 16-bit pixel intensity values within each segmented region (area × mean intensity).

### RNA sequencing of KPC orthotopic allograft tumors from short-term vehicle/morphine-treated mice

RNA sequencing was performed in the Genomics Shared Resource at Roswell Park. Bulk RNA sequencing was performed using the KAPA whole transcriptome library on the Illumina NovaSeq 6000. A volcano plot depicting significantly altered genes was generated in GraphPad Prism 10.6.1 (RRID: SCR_002798). Differential expression analysis was performed on raw gene-level count data using DESeq2 [89] (vDESeq2_1.42.1), which models RNA-seq count data and estimates size factors and dispersions for differential expression testing. For pathway-level analysis, expression data were prepared from DESeq2-normalized counts [90] according to the experimental design and used to perform Gene Set Enrichment Analysis (GSEA) [91] (v4.4.0). GSEA was run against selected MSigDB [92] (v2025) collections, including Hallmark gene sets (H), canonical pathways (C2:CP), and Gene Ontology gene sets [93] (C5:GO). ECM-associated genes were determined through clustering with “ECM organization” in the cancer & cell lines section of The Human Protein Atlas. Enrichr was used to perform pathway analysis using the GO Biological Processes 2025, Kyoto Encyclopedia of Genes and Genomes (KEGG), and Reactome Gene sets [94–96].

### Cell culture

Human immortalized CAF (C7-TA-PSC and PancPat) cells were a gift from Dr. Edna Cukierman (Fox Chase Cancer Center). HTLA cells were a gift from Dr. Brian Roth (University of North Carolina). CAFs were cultured in DMEM (Corning, cat. #MT 10-013-CV) with 10% FBS and 1% P/S added. HTLA cells were cultured in DMEM with 10% FBS and 1% P/S added, as well as 100µg/mL Hygromycin B and 2µg/mL Puromycin. All cells were cultured at 37°C in a 5% CO_2_ incubator.

### PRESTO-Tango protocol

HTLA cells were cultured as indicated above. For transfection, 300,000 HTLA cells/well were plated in a 6-well dish. The next day, Lipofectamine 3000 (Invitrogen, cat. #L3000008) was used according to the manufacturer’s protocol to transfect 1000 ng of either OPRM1-Tango (Addgene, cat. #66464), OPRD1-Tango (Addgene, cat. #66461) or OPRK1-Tango (Addgene, cat. #66462) construct per well. The cells were incubated with the transfection reagent overnight. One well received the transfection reagents without the addition of a construct to serve as a negative control. Following overnight incubation, the cells were replated in a white flat-bottom polystyrene TC-treated 96-well plate (30,000 cells/well). A Beckman Coulter Vi-Cell XR was used to count the cells. On day 4, media was removed from each of the wells and replaced with the respective conditions. On day 5, the Promega Bright-Glo Luciferase Assay System (Promega, cat. #E2610) was used to measure the luminescence of each well according to the manufacturer’s protocol. Extra wells for each condition were plated for the measurement of proliferation via the Promega CellTiter-Glo Luminescent Cell Viability Assay (Promega, cat. #G7571). Average luminescence was normalized to proliferation for each condition, and the resulting fold change in relative receptor activity was calculated.

### Western blot

Protein lysis was performed using the rapid extraction method for mammalian culture outlined by Silva and colleagues [97]. Proteins were separated using 4-15% Mini-Protean TGX precast protein gels (Bio-Rad, cat. #4561083). Proteins were transferred to 0.2µM nitrocellulose membranes (Bio-Rad, cat. #1704270) using the Trans-Blot Turbo Transfer System (Bio-Rad). Membranes were blocked in TBS-T (Tris-buffered saline (TBS) with 0.1% TWEEN-20, Sigma-Aldrich) and 5% w/v bovine serum albumin (BSA) (Fisher Scientific, cat. #BP9700100) for 1 hour at room temperature. Primary antibodies were diluted in 5% BSA in TBS-T and incubated at 4°C overnight. Primary antibodies include the following: COL1A1 (E8F4L) rabbit monoclonal antibody (Cell Signaling Technology, cat. #72026, 1:1000 dilution), COL3A1 (E8D7R) rabbit monoclonal antibody (Cell Signaling Technology, cat. #63034, 1:1000 dilution), αSMA rabbit polyclonal antibody (Proteintech, cat. #14395-1-AP, 1:2000 dilution), and β-actin (8H10d10) mouse monoclonal antibody (Cell Signaling Technology, cat. #3700, 1:1000 dilution). Membranes were washed with TBS-T, and then incubated with horseradish peroxidase-conjugated secondary antibodies (1:2000 donkey anti-rabbit (Fisher Scientific, cat #45-000-682), or 1:2000 goat anti-mouse (Sigma-Aldrich, cat. #A4416)) for 1 hour at room temperature. Pierce ECL Western Blotting Substrate (Thermo Scientific, cat. #32106) was used for chemiluminescent visualization. Signals were developed and imaged using the ChemiDoc XRS+ System and the ChemiDoc Go System with Image Lab Software (Bio-Rad).

### RT-qPCR

Cells were washed once with PBS and once with diH_2_O. RNA was isolated using the Norgen Total RNA Purification Plus Kit (Norgen, cat. #48300) according to the manufacturer’s protocol. RNA concentration and purity were measured via a Thermo Scientific NanoDrop 8000 Spectrophotometer. RNA was stored at -80°C. 900ng of RNA was converted to 20uL of cDNA using the iScript cDNA synthesis kit (Bio-Rad, cat. #1708891) according to the manufacturer’s protocol. cDNA was diluted to 200µL using DEPC water (Invitrogen, cat. #AM9916). qPCR was performed with 10µL reactions using iTaq Universal SYBR Green Supermix (Bio-Rad, cat. #1725121) according to the manufacturer’s protocol using 0.5µL primer and 2µL cDNA per reaction. Thermal cycling was performed using a Bio-Rad CFX Opus 96 Real-Time PCR System. All primers were Bio-Rad PrimePCR SYBR Green Assay primers. Gene expression analysis was performed using the ΔΔCt method.

### siRNA transfection

C7-TA-PSC immortalized human pancreatic CAFs were plated at a density of 200,000 cells/well into a 6-well plate on day 0. On day 1, cells were transfected using Lipofectamine RNAiMAX reagents (Invitrogen, cat. #13778075). The following siRNAs were used: non-targeting control siRNA pool (siCtl) (Horizon Discovery Ltd., cat. #D-001206-13-05), SMARTpool of 4 siRNAs targeting OPRM1 (siOPRM1) (Horizon Discovery Ltd., cat. #L-005686-00-0005), SMARTpool of 4 siRNAs targeting PENK (siPENK) (Horizon Discovery Ltd., cat. #L-020123-00-0005). siRNAs were reconstituted in 1x siRNA buffer (Horizon Discovery Ltd., cat. #B-002000-UB-100) at a stock concentration of 20µM, and the resulting stock solution was aliquoted and stored at -20°C. Transfection was performed by diluting either 1.25µL per well (siOPRM1) or 5µL per well (siPENK) of siRNA into 200µL of Opti-MEM (Gibco, cat. #31985062) and 2µL of RNAiMAX reagent per well. The resulting transfection mixture was briefly vortexed and incubated at room temperature for 20 minutes. During this incubation period, the media on the cells was replaced with 800µL of Opti-MEM. Following the incubation period, 200µL of the transfection mixture was added to each respective well. On day 2, the transfection reagent was removed, and media was replaced with 2mL of DMEM supplemented with 10% FBS and 1% P/S. RNA and protein were collected on day 4 as described above.

### Analysis of single cell RNA sequencing data

Single cell RNAseq data was provided by Loveless et al. [30], which is a previously published atlas where data were aggregated together from prior studies. Briefly, raw data were run through cell ranger pipeline then aligned to the same human reference genome. Data were processed in Seurat V4.4.0 following methods published in Loveless et al. Major cell types were previously defined by expression of lineage specific markers as previously described [98]. Data represent N=26 adjacent normal, N=172 primary tumors, and N=25 metastatic lesions. Tumor samples were obtained from patients with confirmed pancreatic ductal adenocarcinoma. Dot plots are from all patients split by disease state and grouped by cell type. In these plots, the size of the dot represents the relative abundance within each cell type. Blue is adjacent normal, primary tumor is in red, metastases is in black. For DEG, PENK high and PENK low cancer-associated fibroblasts were analyzed. Using Seurat, the CAF population was divided into PENK-positive (expression > 1.5) and PENK-negative (expression ≤ 1.5) groups. The FindMarkers function was then applied to evaluate transcriptomic differences between the two groups.

### Transcriptome-based ECM gene signature and downstream analyses

#### Generation of the MIE gene signature

To derive an extracellular matrix (ECM)-related morphine-induced gene signature, we first analyzed genes upregulated in morphine-treated versus vehicle-treated tumors. Functional enrichment analysis was performed on the morphine-upregulated gene set using Reactome, Gene Ontology (GO), and KEGG pathway databases, with ECM-related pathways consistently identified among the top enriched categories. A candidate 36-gene ECM signature (ECM36) was compiled from morphine-upregulated genes represented in these significantly enriched ECM-related pathways. For downstream clinical and translational analyses, ECM36 was further refined into a 23-gene signature (MIE) by selecting genes based on their consistent prognostic association with patient survival across The Cancer Genome Atlas (TCGA)-based cohorts. Survival associations were evaluated using a publicly available pan-cancer survival analysis resource [99], based on genome-wide Cox proportional hazards models derived from TCGA datasets. Genes demonstrating consistent prognostic significance across multiple cancer types were prioritized for inclusion in the MIE signature. MIE was subsequently used as the principal ECM-related morphine-induced signature in all downstream analyses.

#### Calculation of MIE signature score

MIE signature scores were quantified using a single-sample gene set enrichment analysis (ssGSEA)-based approach. Briefly, normalized gene expression values were used to compute enrichment scores reflecting the coordinated expression of the MIE gene set within individual samples. ssGSEA was performed using the Gene Set Variation Analysis (GSVA) package (version 1.50.5; RRID:SCR_021058) in R (version 4.3.3; RRID:SCR_001905), which applies a rank-based method to estimate per-sample gene set enrichment. All analyses were conducted within individual datasets. The resulting MIE scores were treated as continuous variables for downstream analyses. For survival analyses, MIE scores were dichotomized into high and low groups using the median.

#### Tumor subtype association analysis

To evaluate the association between MIE signature scores and tumor molecular subtypes in pancreatic ductal adenocarcinoma (PDAC), samples from The Cancer Genome Atlas (TCGA)-PAAD cohort were stratified according to three established subtype classification systems: Bailey et al. (progenitor, ADEX, immunogenic, and squamous [28]), Collisson et al. (classical, exocrine-like, and quasi-mesenchymal; [27]), and Moffitt et al. (classical and basal-like; [26]). MIE scores were compared across subtypes within each classification system. For comparisons involving more than two groups (Bailey and Collisson subtypes), statistical significance was assessed using the Kruskal–Wallis test followed by Dunn’s multiple comparisons test. For two-group comparisons (Moffitt subtypes), the Wilcoxon rank-sum test was applied. All analyses were performed within the TCGA-PAAD dataset.

#### Survival analysis

The prognostic significance of the MIE signature was evaluated across multiple cancer types using TCGA pan-cancer dataset. Overall survival (OS) and, where applicable, disease-free survival (DFS) were used as clinical endpoints. Patients were stratified into MIE-high and MIE-low groups based on median signature scores. Kaplan–Meier survival curves were generated to estimate survival distributions, and statistical significance between groups was assessed using the log-rank (Mantel–Cox) test. Hazard ratios (HRs) were estimated using Cox proportional hazards regression models where applicable. Pan-cancer survival analyses were conducted using the GEPIA2 web server (Gene Expression Profiling Interactive Analysis 2; [100]), a publicly available platform based on uniformly processed TCGA RNA-seq data, enabling survival analysis through gene expression–based stratification.

#### Correlation analysis

Correlation analyses were performed to evaluate the relationship between MIE scores and other transcriptional or phenotypic features. Spearman rank correlation coefficients were calculated.

## Statistical Analysis

Statistical analyses were performed in GraphPad Prism 10.6.1 (RRID: SCR_002798). Asterisks on graphs indicate statistically significant differences: *, *p* < 0.05; **, *p* < 0.01; ***, *p* < 0.001; **** *p* < 0.0001.

---

**Supplementary Figure 1.**
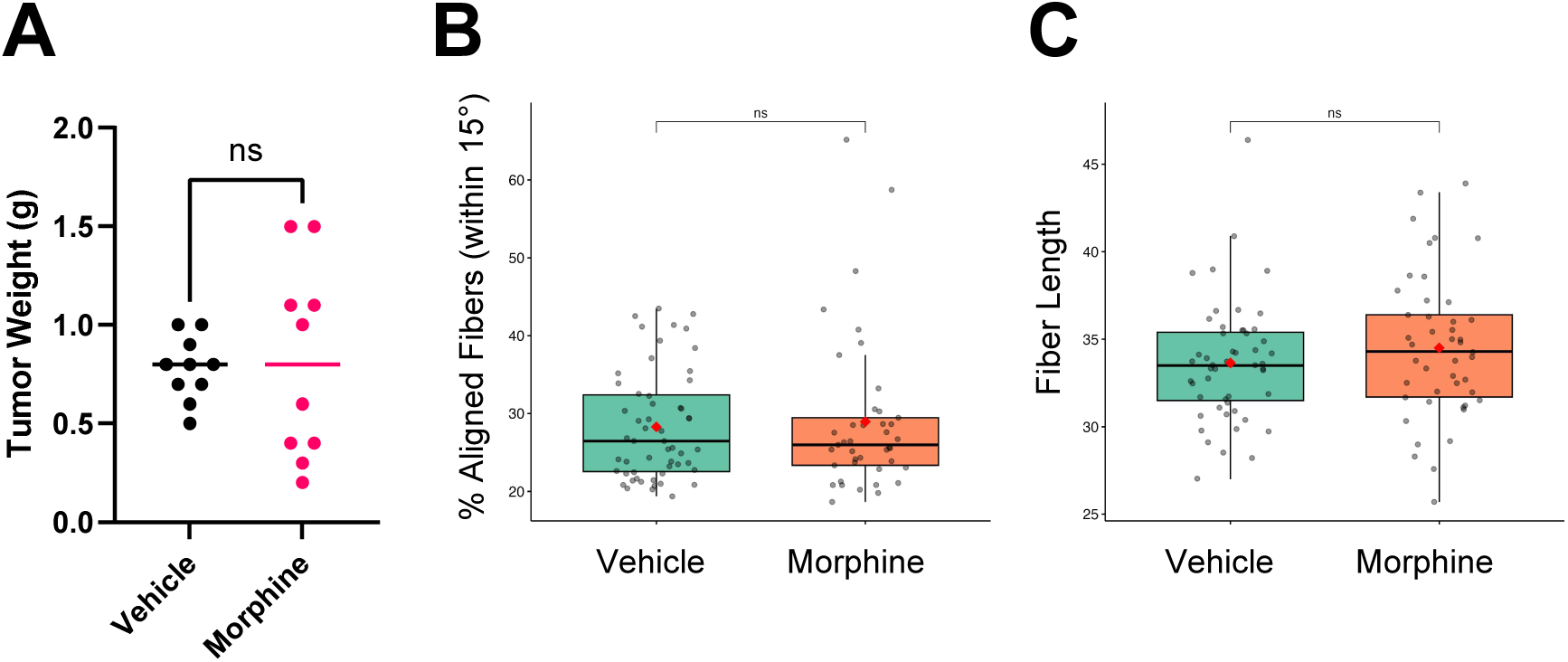
: Morphine does not significantly alter pancreatic tumor weight or collagen fiber alignment. **A,** Weight of tumors (g) from mice treated with either vehicle or morphine, *n* = 10 for each arm. An unpaired T test was used for analysis of statistical significance. **B,** Quantification of collagen fiber alignment assessed by SHG microscopy in tumors from mice treated with either vehicle or morphine. **C,** Quantification of collagen fiber length assessed by SHG microscopy in tumors from mice treated with vehicle or morphine. Statistical analyses in **B** and **C** were performed using the Wilcoxon test.

**Supplementary Figure 2.**
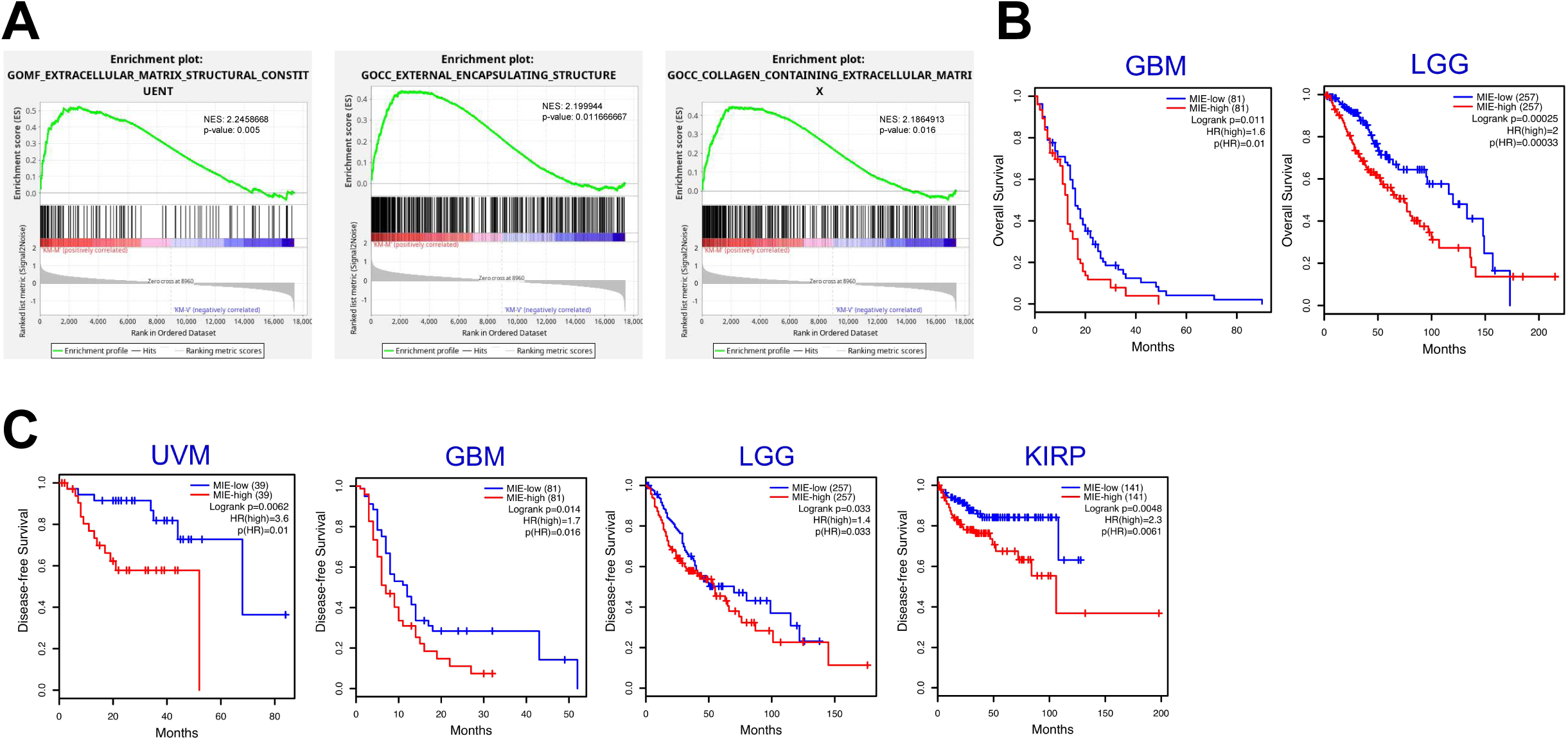
: Morphine induces expression of an extracellular matrix signature that correlates with worsened overall and disease-free survival across multiple cancer types outside of PDAC. **A,** Enrichment plots of GOMF_Extracellular_Matrix_Structural_Constituent (*p* = 0.005), GOCC_External_Encapsulating_Structure (*p =* 0.011666667), and GOCC_Collagen_Containing_Extracellular_Matrix (*p* = 0.016). **B,** Elevated MIE predicts poor overall survival (OS) in glioblastoma (GBM) and low grade glioma (LGG) based on TCGA survival analyses. **C,** Elevated MIE predicts poor disease-free survival (DFS) in uveal melanoma (UVM), GBM, LGG, and kidney renal papillary cell carcinoma (KIRP) based on TCGA survival analyses. Kaplan-Meier survival curves were used to estimate survival distributions, and statistical significance between groups was assessed using the log-rank (Mantel-Cox) test. Hazard ratios (HRs) were estimated using Cox proportional hazards regression models. *, *p* < 0.05; **, *p* < 0.01; ***, *p* < 0.001; **** *p* < 0.0001. Sample sizes are indicated in parenthesis.

**Supplementary Figure 3.**
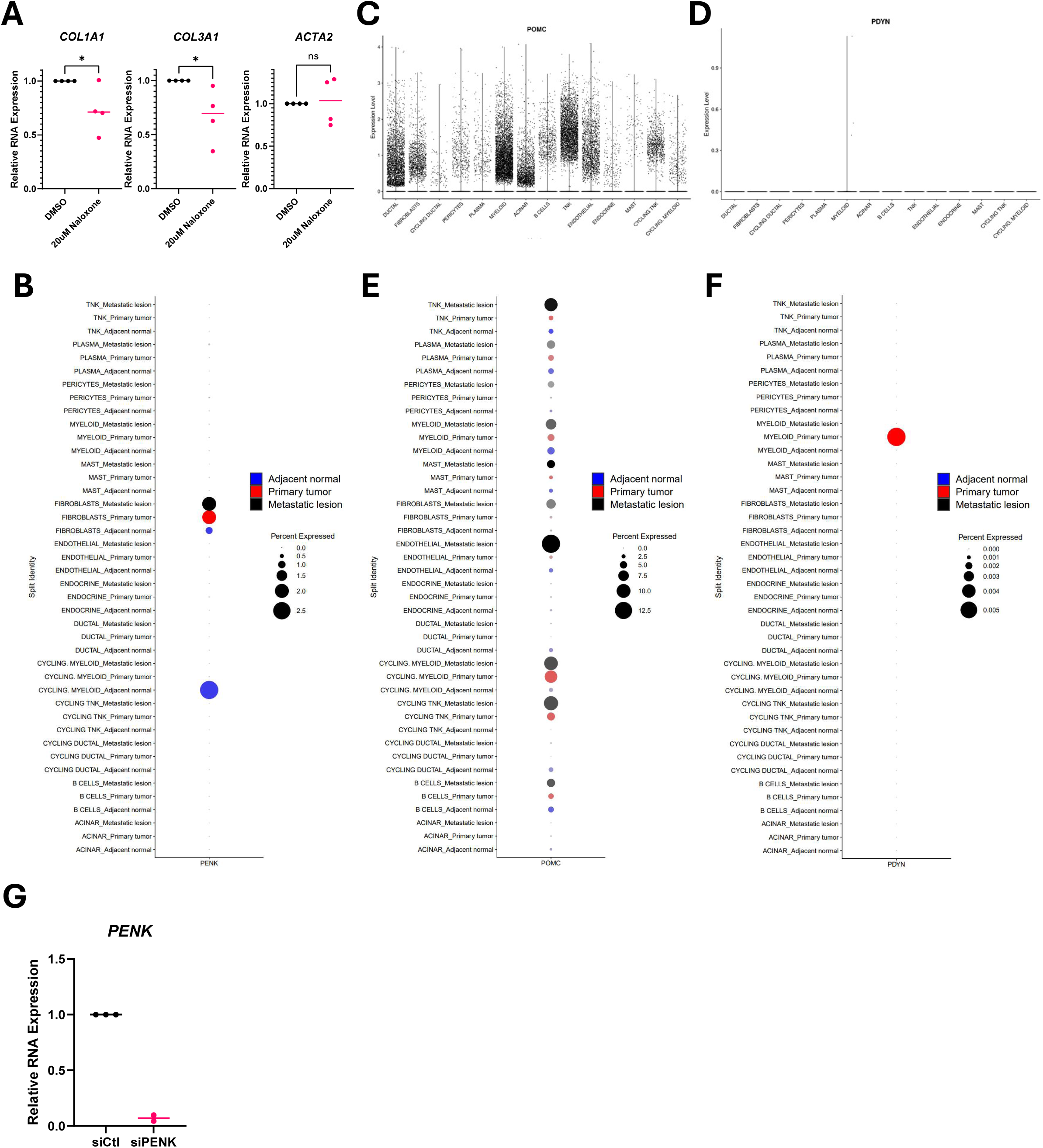
: PENK is the dominant endogenous opioid precursor expressed in CAFs. **A,** qPCR for *COL1A1*, *COL3A1*, and *ACTA2* of immortalized human PancPat pancreatic CAFs treated with 20µM vehicle or naloxone for 6 hours. Unpaired T tests were used for the assessment of statistical significance between groups. **B,** Dot plot depicting expression of PENK across cell types in adjacent normal tissue, metastatic lesions, and primary tumors using human single cell RNA sequencing data. **C,** Expression of POMC by cell type in primary tumors using human single cell RNA sequencing data. **D,** Expression of PDYN by cell type in primary tumors using human single cell RNA sequencing data. **E,** Dot plot depicting expression of POMC across cell types in adjacent normal tissue, metastatic lesions, and primary tumors using human single cell RNA sequencing data. **F,** Dot plot depicting expression of PDYN across cell types in adjacent normal tissue, metastatic lesions, and primary tumors using human single cell RNA sequencing data. **G,** qPCR for *PENK* in siCtl or siPENK immortalized human C7-TA-PSC pancreatic CAFs. *, *p* < 0.05; **, *p* < 0.01; ***, *p* < 0.001; **** *p* < 0.0001.

**Supplementary Figure 4.**
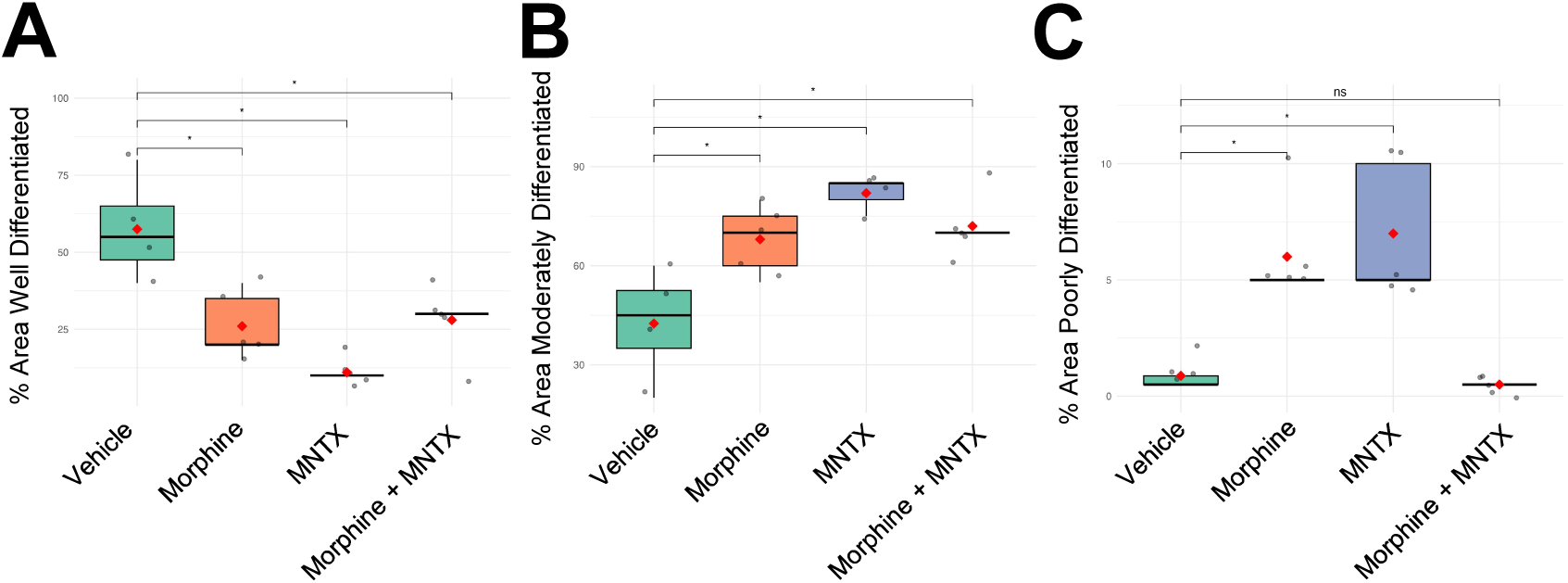
: MNTX alters the differentiation status of pancreatic tumors. **A,** Quantification of well differentiated areas of tumors from mice treated with vehicle (*n* = 4), morphine (*n* = 5), MNTX (*n* = 5), or morphine + MNTX (*n* = 5). **B,** Quantification of moderately differentiated areas of tumors from mice treated with vehicle, morphine, MNTX, or morphine + MNTX. **C,** Quantification of poorly differentiated areas of tumors from mice treated with vehicle, morphine, MNTX, or morphine + MNTX. Comparisons were made for assessment of statistical significance using the Wilcoxon test. *, *p* < 0.05; **, *p* < 0.01; ***, *p* < 0.001; **** *p* < 0.0001.

